# Label-free biosensor assay decodes the dynamics of Toll-like receptor signaling

**DOI:** 10.1101/2024.01.18.576268

**Authors:** Janine Holze, Felicitas Lauber, Evi Kostenis, Günther Weindl

**Affiliations:** Pharmacology and Toxicology Section, Pharmaceutical Institute, University of Bonn, 53121, Bonn, Germany; Molecular, Cellular and Pharmacobiology Section, Institute of Pharmaceutical Biology, University of Bonn, 53115, Bonn, Germany

## Abstract

The discovery of Toll-like receptors (TLRs) represented a significant breakthrough that paved the way for the study of hostpathogen interactions in innate immunity. However, there are still major gaps in understanding TLR function, especially the early dynamics of downstream TLR pathways remains less clear. Here, we present a label-free optical biosensor-based assay as a powerful method for detecting TLR activation in a native and label-free environment and defining the dynamics of TLR pathway activation. This technology is sufficiently sensitive to detect TLR signaling and readily discriminates between different TLR signaling pathways. We define pharmacological modulators of cell surface and endosomal TLRs and downstream signaling molecules and uncover previously unrecognized TLR signaling signatures, including biased receptor signaling. These findings highlight that optical biosensor assays complement traditional assays that use a single endpoint and have the potential to facilitate the future design of selective drugs targeting TLRs and their downstream effector cascades.

## Introduction

Toll-like receptors (TLRs) form the first barrier in the innate immune response and therefore represent a highly interesting and therapeutically promising target to modulate immune response and inflammation^1,2^. TLRs detect pathogen-associated molecular patterns (PAMPs) of invading microorganisms that enable the host to distinguish between different infections. In addition to PAMPs, also stressor injuryreleased molecules, termed damageor danger-associated molecular patterns (DAMPs), can also be sensed by TLRs. Mechanistically, TLRs dimerize for activation, and trigger intracellular pathways that promote diverse cellular responses, including inflammatory processes^3^. Most of our knowledge on TLR signaling comes from loss-of-function genetic analysis and TLR modulation by small-molecule ligands and has been studied mainly by analyzing a limited set of downstream signaling pathways and effector proteins such as cytokines. Likewise, fluorescent tags or reporter systems that are prone to non-specific interference have been used to study TLR function and signaling^4^. Given these limitations, new approaches that allow a holistic view on TLR signaling and its modulation are highly desired^5^.

In G protein coupled receptor (GPCR) research, the concept of analyzing whole cell responses – rather than individual components of signaling pathways – has been well established^6^. The approach using label-free whole-cell biosensing, for example based on dynamic mass redistribution (DMR), not only offers access to rich cellular information but also provides new and exciting mechanistic insights into GPCR activation and ligand-dependent modulation^7–9^. This technology uses a polarized broadband light that passes through the bottom of a biosensor microtiter plate containing the cells. Upon receptor activation, a shift in the wavelength of the reflected light reveals signal-transduction events linked to changes in cell morphology^10^. Depending on the signaling pathways activated and the cell-shaped change-related mass redistribution, the direction (positive or negative) and magnitude of the detected shift in wavelength may vary (Fig. 1a). Optical biosensor assays have the main advantage in analyzing living cells in real time and identifying signal events by detecting whole-cell responses related to changes in cellular shape^11^. These morphological changes can be observed as long as the signaling event results in a rearrangement of the cytoskeleton^6,12^. Unlike traditional assays that use a single endpoint, DMR offers the possibility of detecting the cumulative response of living cells by high-throughput screening. This opens up new possibilities for exploring and defining pathway-selective effector activation or biased signaling of TLRs. The latter represents the ability of a ligand to selectively activate some but not all signaling pathways and is a well-established paradigm for GPCRs^13,14^. Other receptor classes, such as TLRs, also trigger multiple signaling pathways, but their bias signaling capabilities have not yet been established^4,15^.

**Figure 1:**
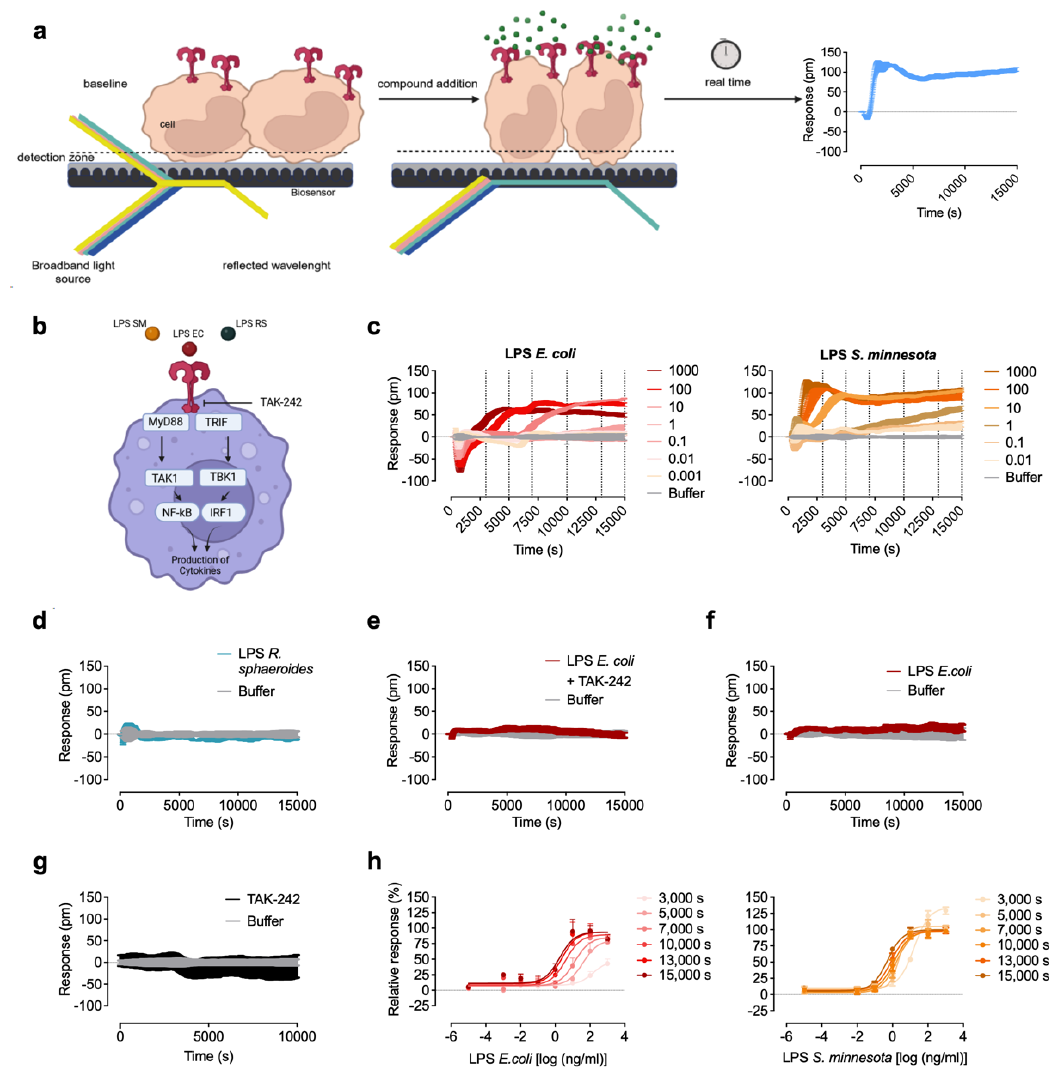
Optical biosensor discriminates different LPS chemotypes in HEK293 TLR4 reporter cells. **(a)** DMR principle: cultured cells are grown in 384 well microplates equipped with a resonant waveguide grating biosensor within the bottom of each well. A specific broadband light source illuminates the lower surface of the biosensor. The illumination generates an energy field within the DMR detection zone, and its strength diminishes exponentially with the distance from the sensor. Under baseline conditions (left) cells are in close proximity to the biosensor, and the refractive index of these cells determines the reflected wavelength, which is subsequently measured. If cells undergo morphological changes (middle), the refractive indices above the sensor can either increase or decrease. Consequently, the reflected wavelength becomes shorter or longer, respectively. The shift in the measured wavelength is plotted against the recording time (right). **(b)** Schematic representation of TLR4 signaling and contact point of TAK-242. **(c,d)** Baseline corrected DMR representative recordings of HEK293 TLR4 reporter cells stimulated with the indicated concentrations (ng/ml) of LPS from *E. coli, S. minnesota* and *R. sphaeroides*. **(e,f)** Baseline corrected DMR representative recordings of HEK293 TLR4 reporter cells preincubated with 50 *μ*M of the TLR4-antagonist TAK-242 (e) or HEK293 control reporter cells lacking TLR4 **(f)** stimulated with LPS *E. coli* (1 *μ*g/ml). **(g)** Baseline corrected DMR representative recordings of HEK293 TLR4 reporter cells treated with TAK-242 (50 *μ*M). Shown are representative data of at least three independent experiments. **(h)** Sigmoidal concentration effect curves resulting from DMR traces **(c)** of at least three independent experiments (Mean ± S.E.M). Concentration-effect curves of DMR data were generated by the response at six different time points. Calculated pharmacological parameters of the concentration-effect curves are depicted in table 1.

The chemical toolbox for studying the dynamics of TLR signaling has steadily increased^16^, however, label-free approaches are still very limited. Here, we demonstrate that label-free detection captures receptor activation and signaling of TLRs in real time, which has not been reported by other technology platforms. We reveal unrecognized mechanisms of TLR pathway activation and biased signaling, thus providing a completely new insight into TLR signal transduction.

**Table 1:**
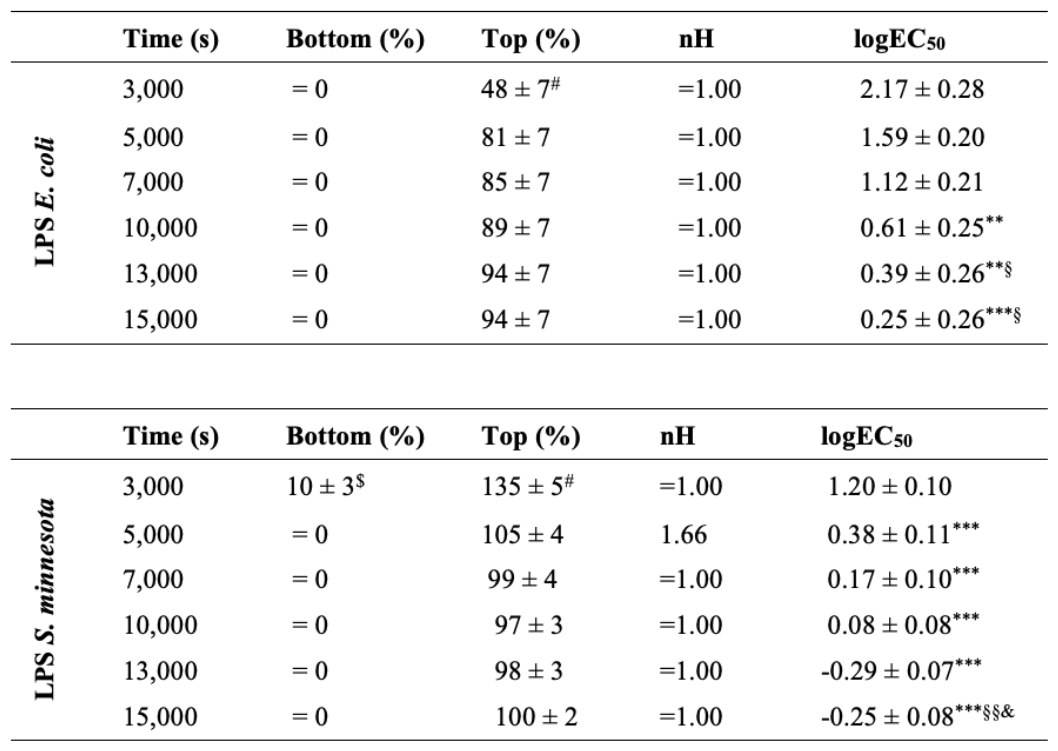
Pharmacological parameter of LPS induced DMR at six selected time points in HEK293 TLR4 reporter cells. Curve fitting of single experiments was obtained by nonlinear regression analysis applying a three (nH = 1.00) or four-parameter logistic equation. Values are means ± S.E.M. of at least three independent experiments. ^#^P<0.05 significantly different from 100%, ^$^P<0.05 significantly different from 0% (one-sample t-test). **P<0.01, ***P<0.001 significantly different from values at 3,000 s. ^§^P<0.05, ^§§^P<0.01 significantly different from values at 5,000 s. ^&^P<0.05 significantly different from values at 7000 s, one-way analysis of variance (ANOVA, Tukey’s post-test).

## Results

### Label-free optical biosensor assay discriminates TLR4 signaling of LPS chemotypes

TLR4 is activated by lipopolysaccharide (LPS), a key component of the outer membrane of Gram-negative bacteria^17^. LPS, consists of lipid A, a core oligosaccharide, and an O side chain^18^. In particular, the hydrophobic component, lipid A, is responsible for its immune stimulating activity. The structures of lipid A in various bacteria display variability in the number or length of attached fatty acids, a critical factor for TLR4 activation. This activation of the TLR4 subtype by LPS is known to initiate the MyD88 signaling pathway at the plasma membrane, as well as CD14-dependent endocytosis, which initiates the TRIF-dependent cascade^19–21^ (Fig. 1b).

Currently, it is known that LPS from various sources elicits distinct inflammatory responses and exhibits a form of biased signaling^4^. To test whether label-free cell-based assays using optical biosensor technology can be reliably used to study TLR signaling and discover potentially biased signaling, we stimulated HEK293 reporter cells, stably transfected to express TLR4, with increasing concentrations of the TLR4 ligand LPS *Escherichia coli* (LPS EC) and DMR signals were recorded (Fig. 1c). No signals were detected in the presence of the TLR4 antagonist TAK-242 (Fig. 1e) or control HEK293 cells lacking TLR4 (Fig. 1f), demonstrating that the LPS-induced DMR signal originates specifically from TLR4 activation. The TLR4 inhibitor TAK-242 alone induced no detectable signal (Fig. 1g). HEK293 TLR4 and control reporter cells lacking TLR4 (HEK293 Null2) showed characteristic DMR signals of G protein signaling^6,22^ in the presence of the GPCR ligands acetylcholine, epinephrine, or atropine, respectively. This illustrates that cells overexpressing TLR4 show consistent GPCR signaling patterns compared to control cells lacking TLR4 (Supplementary Figs. S1a and b). LPS from *E. coli* concentration-dependently induced an early negative peak at 720 s followed by a large continuing positive change in DMR signals for concentrations above 10 ng/ml (Fig. 1c). The DMR response distinctly varied for LPS from *S. minnesota*, displaying an early and robust positive signal starting at 1,500 s. For both LPS chemotypes, positive signals did not return to baseline during the 4 h recording period. In contrast, LPS from *Rhodobacter spharoides* (LPS RS), a TLR4 antagonist^23^, induced no detectable signal (Fig. 1d).

To gain a deeper understanding of the pharmacology related to DMR signals induced by the different LPS and to study the kinetics of receptor potency (EC_50_) and efficacy (E_max_) at early signaling events, we established full concentration-effect curves generated by using six different time points (Fig. 1h). We found a time-dependent increase in potency and efficacy for LPS from *E. coli* that remained constant after 13,000 s (Table 1). The efficacy increased after 3,000 s and remained unchanged throughout the recording time. For LPS from *S. minnesota* we observed a similar time-dependent change in potency, whereas the efficacy decreased after 5,000 s. Collectively, these data indicate that DMR readily discriminates between LPS of different bacteria and uncovers a differential concentrationand chemotypedependent signaling signature.

### Optical biosensor assay uncovers differential signaling of TLR2 heterodimers

TLR2 operates as a heterodimer with TLR1 or TLR6^24^. TLR2/1 and TLR2/6 heterodimers are known to initiate similar signaling cascades, resulting in comparable gene activation profiles^25^. However, we and others have reported variations in signaling and sensitivity to ligands for different TLR2 heterodimers^26–31^. Furthermore, agonists for TLR2/1 and TLR2/6 induce NF-κB, MAP kinase activation, and cytokine transcription with different kinetics^32^. To assess whether TLR2 activation and signal variances are also detectable with whole-cell biosensing, we focused on TLR2 (Fig. 2a). To ensure specific activation of the receptor, we used the agonists Pam_3_CSK_4_ (TLR2/1) and Pam_2_CSK_4_ (TLR2/6), and HEK293 reporter cells that are stably transfected with only one of the heterodimers after knockout of endogenous TLR1 or TLR6, respectively^31^. In line with our previous observations, Pam_3_CSK_4_ predominantly activated TLR2/1, while Pam_2_CSK_4_ mainly activated TLR2/6 in a concentration-dependent manner (Fig. 2b and Supplementary Fig. 2a). No signals were detected in TLR4 reporter cells (Supplementary Fig. 2b) or control HEK293 cells lacking TLR1 and 6 (Supplementary Fig. 2c). The latter cells also showed characteristic DMR signals of G protein signaling in the presence of acetylcholine or epinephrine, respectively (Supplementary Fig. 2d). Similar to LPS, stimulation with Pam_3_CSK_4_ and Pam_2_CSK_4_ resulted in a positive DMR signal that persisted throughout the 4 h recording period. Both TLR2 agonists showed distinct DMR profiles, with Pam_3_CSK_4_ inducing a faster early decline followed by an increase compared to Pam_2_CSK_4_. The concentration-effect curves (Fig. 2c) revealed a different kinetic dependence compared to LPS. The efficacy of Pam_3_CSK_4_ decreased more pronounced and gradually over time (Table 2), while its potency remained constant. In comparison, the TLR2/6 agonist Pam_2_CSK_4_ demonstrated constant efficacy, but the potency decreased significantly after 13,000 s. These findings offer evidence to suggest that both agonists trigger differences in the recruitment of adaptor proteins, including MyD88 (Myeloid differentiation primary response 88) for the MyD88-dependent signaling pathway and Mal (Myd88 adaptor-like protein) for the MyD88independent pathway^32^ as well as signal cascade kinetics.

**Table 2:** Pharmacological parameters of Pam_3_CSK_4_and Pam_2_CSK_4_-induced DMR at six selected time points in HEK293 TLR2/1 and TLR2/6 reporter cells. Curve fitting of single experiments was obtained by nonlinear regression analysis applying a three (nH = 1.00) parameter logistic equation. Values are Means ± S.E.M. of at least three independent experiments. ^#^P<0.05, ^##^P<0.01 significantly different from 100% (one-sample t-test). *P<0.05, **P<0.01 significantly different from values at 3,000 s. ^§^P<0.05 significantly different from values at 5,000 s, one-way analysis of variance (ANOVA, Tukey’s post-test).

|  | Time (s) | Bottom (%) | Top (%) | nH | logEC<sub>50</sub> |
| --- | --- | --- | --- | --- | --- |
| TLR2/1<br>Pam<sub>3</sub>CSK<sub>4</sub> | 3,000 | = 0 | 163 ± 11<sup>##</sup> | =1.00 | 1.80 ± 0.12 |
|  | 5,000 | = 0 | 152 ± 9<sup>##</sup> | =1.00 | 1.72 ± 0.11 |
|  | 7,000 | = 0 | 128 ± 8<sup>#</sup> | 2.17 ± 0.76 | 1.42 ± 0.16 |
|  | 10,000 | = 0 | 140 ± 12<sup>#</sup> | =1.00 | 1.44 ± 0.18 |
|  | 13,000 | = 0 | 118 ± 12 | =1.00 | 1.30 ± 0.22 |
|  | 15,000 | = 0 | 96 ± 11<sup>**§</sup> | =1.00 | 1.20 ± 0.26 |
| TLR2/6<br>Pam<sub>2</sub>CSK<sub>4</sub> | 3,000 | = 0 | 88 ± 14 | =1.00 | 0.39 ± 0.34 |
|  | 5,000 | = 0 | 105 ± 12 | =1.00 | -0.36 ± 0.30 |
|  | 7,000 | = 0 | 101 ± 11 | =1.00 | -0.73 ± 0.29 |
|  | 10,000 | = 0 | 96 ± 10 | =1.00 | -0.90 ± 0.27 |
|  | 13,000 | = 0 | 97 ± 7 | =1.00 | -1.05 ± 0.21* |
|  | 15,000 | = 0 | 95 ± 6 | =1.00 | -1.08 ± 0.19* |

**Figure 2:**
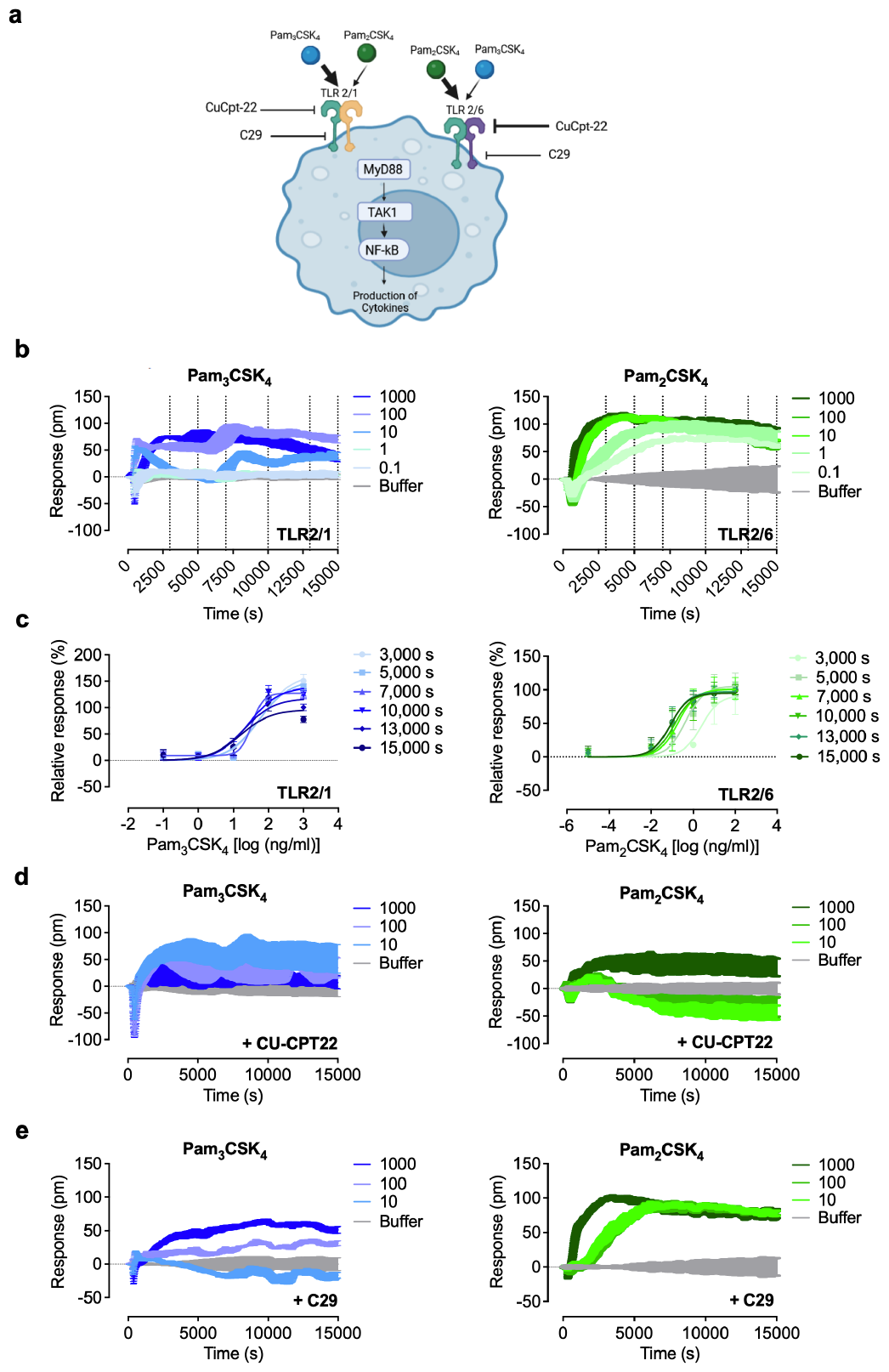
Optical biosensor assays detect differential signaling of TLR2 heterodimers. **(a)** Schematic representation of TLR2 signaling, Ligand priority and contact point of the two antagonists CU-CPT22 and C29. **(b)** Baseline corrected DMR representative recordings of HEK293 TLR2/1 or 2/6 reporter cells stimulated with the indicated concentrations (ng/ml) of Pam_3_CSK_4_ or Pam_2_CSK_4_. **(c)** Sigmoidal concentration-effect curves resulting from DMR traces of at least three independent experiments (Mean ± S.E.M). Concentration-effect curves of DMR data were generated by the response at six different time points. **(d,e)** Baseline corrected DMR representative recordings of HEK293 TLR2/1 or 2/6 reporter cells preincubated with 50 *μ*M of the TLR2 antagonist CU-CPT22 **(d)** or C29 **(e)** stimulated with the indicated concentrations (ng/ml) of Pam_3_CSK_4_ or Pam_2_CSK_4_. Shown are representative data of at least three independent experiments. Calculated pharmacological parameters of the concentration-effect curves **(c)** are depicted in table 2.

Next, we tested two TLR2 antagonists which differ in inhibitory potency and heterodimer predominance^44^ (Fig. 2a). We confirmed that CU-CPT22 preferentially inhibits Pam2CSK4 signaling, whereas C29 shows only weak effects for both heterodimers (Figs. 2a, d and e). Together, our data demonstrate the detectability of TLR2 and subsequent pathway activation. Furthermore, we have shown that heterodimerization of TLR2 with TLR1 or TLR6 leads to clearly distinct optical signals, indicating differential signaling behavior.

### Optical biosensor assay detects TLR subtypes in native cell lines

Label-free biosensor assays offer the possibility of investigating receptor activation in a native cell setting^33^. We tested whether the technology can be applied to cell lines of different origin that express endogenous TLRs. We used the well-characterized epidermal HaCaT cell line that responds to TLR2/1 and TLR2/6 ligands, but shows no functional TLR4 signaling^34^. As expected, stimulation with LPS from *E. coli* generated a signal identical to control, while Pam_3_CSK_4_ and Pam_2_CSK_4_ concentration-dependently induced DMR responses (Fig. 3a and Supplementary Fig. 3a). The signal generated by Pam_3_CSK_4_ was nearly indistinguishable from that seen in HEK293 cells expressing TLR2/1 (Figs. 2b and 3b). However, the inherent cellular phenotype should be considered when interpreting the data obtained in both cell lines. The DMR signal for Pam_2_CSK_4_ was different in HaCaT cells and TLR2/6 reporter cells. HaCaT cells lack the expression of TLR6^34^ and Pam_2_CSK_4_ can activate TLR2/1 heterodimers^31,35^, therefore we reasoned that the TLR2/6 ligand triggers TLR2/1 in HaCaT cells. Indeed, the signal for Pam_2_CSK_4_ obtained in TLR2/1 reporter cells was close to that detected in HaCaT cells and clearly differed from Pam_3_CSK_4_ (Fig. 2b and Supplementary Fig. 2a). In the presence of the TLR2 antagonists CU-CPT22 and C-29, the Pam_3_CSK_4_-induced responses were blunted (Fig. 3c). Both antagonists induced initial transient signals that returned to baseline before stimulation with Pam_3_CSK_4_ (Supplementary Fig. 3b), thus, overlapping signals can be excluded. The optical traces triggered by the lipopeptides were independent of TLR4 (Supplementary Fig. 3c). For control purposes, we confirmed that acetylcholine and epinephrine, which act via endogenous GPCRs, and the adenylyl cyclase-activating agent forskolin showed the signal trace fingerprints typical for HaCaT cells^36^ (Supplementary Fig. 3d).

**Figure 3:**
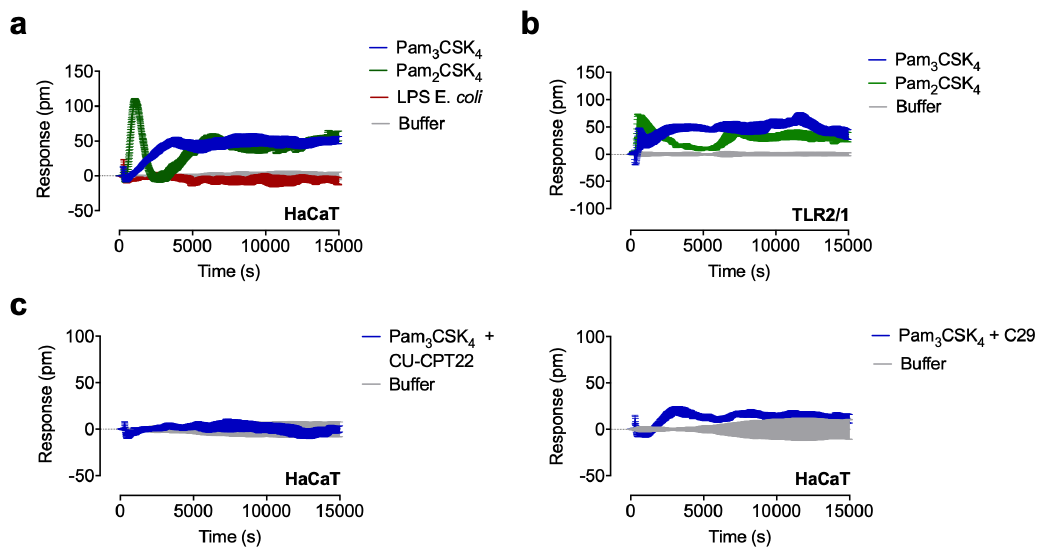
Optical biosensor uncovers TLR2 and TLR4 signaling in epidermal HaCaT cells. **(a)** Baseline corrected DMR representative recordings of HaCaT cells stimulated with LPS *E. coli* (1000 ng/ml), Pam_3_CSK_4_ (1000 ng/ml) or Pam_2_CSK_4_ (1000 ng/ml). **(b)** Baseline corrected DMR representative recordings of HEK293 TLR2/1 reporter cells stimulated with Pam_3_CSK_4_ (100 ng/ml) or Pam_2_CSK_4_ (100 ng/ml). **(c)** Baseline corrected DMR representative recordings of HaCaT cells preincubated with 50 *μ*M of the TLR2 antagonists CU-CPT22 or C29 treated with Pam_3_CSK_4_ (1000 ng/ml). Shown are representative data of at least three independent experiments.

### Label-free optical biosensor assay reveals TLR ligand bias

Next, we tested whether TLR signaling can be detected in THP-1 cells, as DMR can be performed with cells expressing endogenous level of receptor as well as with suspension cells^6^. The monocytic cell line grows in suspension but can be induced to differentiate into adherent macrophage-like cells. In line with the poor response of THP-1 monocytes to LPS from E. coli^37^, only a weak negative signal was observed in undifferentiated THP-1 cells (Fig. 4a) while PMA-differentiated cells responded to *E. coli* LPS with robust signals (Fig. 4b). LPS from *S. minnesota*, on the other hand, could elicit a detectable signal in both THP-1 monocytes and macrophages (Figs. 4a and b). The different ability to induce strong signals in THP-1 monocytes can also be ascribed to different characteristics in LPS. LPS from *E. coli* is a smooth (s)-form LPS whereas LPS from *S. minnesota* is assigned to the rough (r)-form^38^. Both share the same receptor complex, but their mechanism of action differs. Although CD14 is essential for sLPS-induced NF-KB und IRF signaling, its presence is not required for the rLPS-induced pathway^39^. Since THP-1 monocytes express low levels of CD14^40^, these observations can be confirmed by our results.

**Figure 4:**
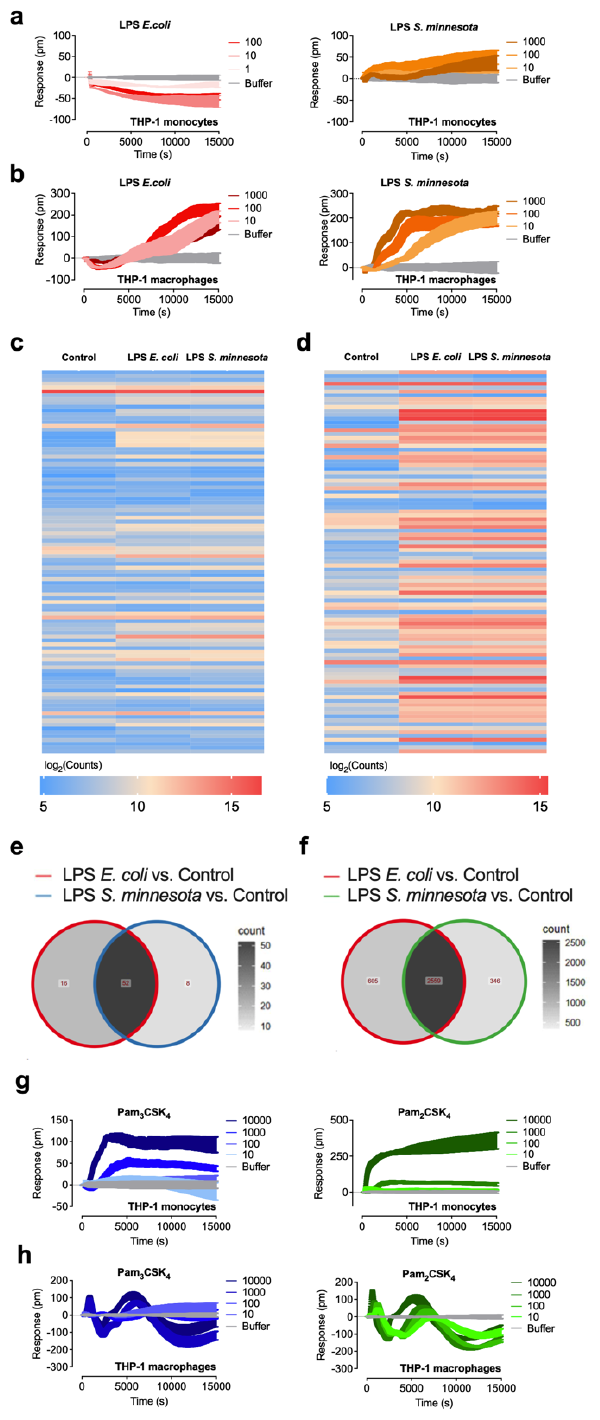
Optical biosensor reveals differences in TLR2 signaling in THP-1 monocytes and macrophages. **(a,b)** Baseline corrected DMR representative recordings of THP1 monocytes **(a)** or macrophages **(b)** stimulated with increasing concentrations (ng/ml) of LPS *E*.*coli* and LPS *S. minnesota*. **(c,d)** Heatmap of the top 100 significant upor downregulated genes identified in HEK293-TLR4 cells **(c)** or THP-1 macrophages **(d)** treated with buffer only (control), LPS *E*.*coli* or LPS *S. minnesota*, after 3 h incubation. The global transcriptional response induced by the LPS chemotypes showed no significant differences. **(e,f)** Venn diagram for HEK293-TLR4 cells **(e)** and THP-1 macrophages **(f)** indicating the number of significant (FDR<0.05) differentially expressed genes and the overlap between each set of genes treated with buffer only (control), LPS *E. coli* or LPS *S. minnesota*, after 3 h incubation. **(g,h)** Baseline corrected DMR representative recordings of THP-1 monocytes **(g)** or macrophages **(h)** stimulated with increasing concentrations (ng/ml) Pam_2_CSK_4_ or Pam_3_CSK_4_. Shown are representative data of at least three independent experiments.

Both LPS chemotypes induce unique signal fingerprints at early time points. Thus, we analyzed whether the different signals lead to differential transcriptional responses. We performed RNA-seq of TLR4-HEK293 cells and THP-1 macrophages after 3 h stimulation with LPS from *E. coli* and *S. minnesota*, respectively. In both cell lines, the global transcriptional response induced by the LPS chemotypes was similar (Figs. 4c and d). As expected, in TLR4-HEK293 cells, the LPS chemotypes induced a weak transcriptional response with 52 genes differentially expressed compared to control (Fig. 4e), whereas a strong response with 2559 differentially expressed genes was observed in THP-1 macrophages (FDR adjusted p-value <0.05 and |logFC | > 1) (Fig. 4f). *E. coli* LPS specifically modulated 16 and 605 genes in TLR4-HEK293 cells and THP-1 macrophages, respectively. In contrast, around two-fold less genes were specifically regulated by LPS from *S. minnesota*. However, the volcano plots showed no significant differences between both LPS chemotypes, except for one gene in HEK293 TLR4 cells (Supplementary Figs. 4a and b).

We next stimulated THP-1 monocytes with the TLR2 ligands Pam_3_CSK_4_ and Pam_2_CSK_4_ and found early positive signals that increased over time reaching a plateau at 2,700 and 1,800 sec, respectively (Fig. 4g). We observed a marked difference in the optical trace signatures in THP-1 macrophages (Fig. 4h). The signals for Pam_3_CSK_4_ and Pam_2_CSK_4_ clearly resembled oscillatory dynamics with a fast initial peak at 720 and 600 s, a negative peak at 2,100 and 1,800 s followed by a second positive peak at 5,100 and 4,800 s before finally reaching a second negative peak at 10,800 and 10,500 s, respectively.

Using HEK293 cells expressing specific TLRs, as well as native cell lines such as HaCaT cells and THP-1 monocytes and macrophages, we have demonstrated that agonist-induced signals can be differentiated by label-free technology.

Next, we sought to study TLR dynamics in primary monocytes isolated from peripheral blood mononuclear cells (PBMCs). As expected, nontoxic concentrations of the tested TLR4 and TLR2 agonists induced cytokine secretion (Supplementary Figs. 5a and b). LPS from *E. coli* and *S. minnesota* displayed concentration-dependent positive signals (Fig. 5a) which were inhibited in the presence of TAK-242 (Fig. 5b). In line with the results in cell lines, *S. minnesota* LPS showed a faster signal increase compared to *E. coli* LPS. The TLR2/6 agonist Pam_2_CSK_4_ induced concentration-dependent signals, whereas the TLR2/1 agonist Pam_3_CSK_4_ showed signals only at the highest concentration (Fig. 5c). Our findings demonstrate that TLR2 and TLR4 signaling differs between cell types and shows unique optical signatures.

**Figure 5:**
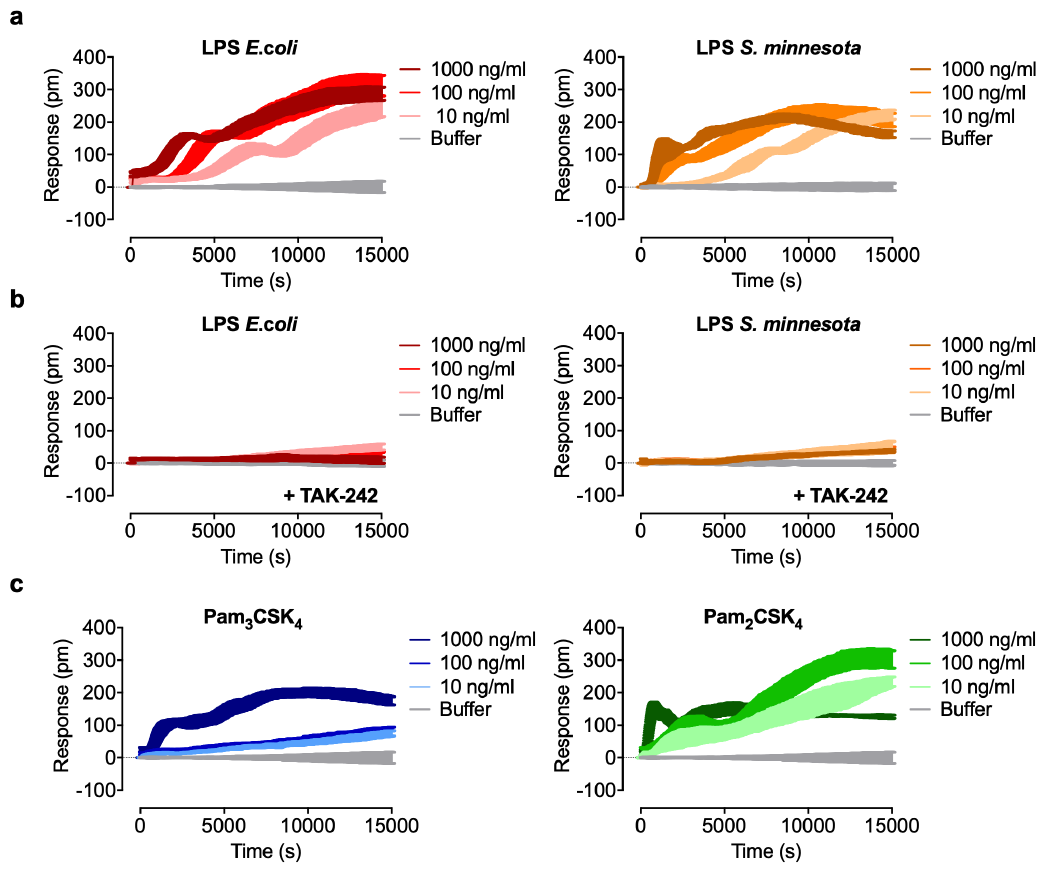
Optical biosensor assay decodes signaling in primary monocytes. **(a)** Baseline corrected DMR representative recordings of primary monocytes isolated from PBMCs stimulated with the indicated concentrations (ng/ml) of LPS from *E. coli* or *S. minnesota*. **(b)** Baseline corrected DMR representative recordings of primary monocytes isolated from PBMCs preincubated with 50 *μ*M of the TLR4-antagonist TAK-242 and stimulated with the indicated concentrations (ng/ml) of LPS from *E. coli* or *S. minnesota*. **(c)** Baseline corrected DMR representative recordings of primary monocytes isolated from PBMCs stimulated with the indicated concentrations (ng/ml) of Pam_3_CSK_4_ and Pam_2_CSK_4_. Shown are representative data of at least three independent experiments and donors.

### LPS chemotypes differentially induce MyD88-dependent signaling

TLR4 induces both MyD88-dependent and -independent pathways^41,42^. MyD88 triggers rapid NF-ΚB activation and the release of proinflammatory cytokines such as tumor necrosis factor-alpha (TNF), interleukin (IL-)1β, IL-6 and IL-8. The MyD88-independent pathway leads to rapid activation of interferon regulatory factor (IRF)3, resulting in the release of interferon (IFN)-β and delayed NF-ΚB activation^43^. Based on the differences identified by the optical biosensor in the early response to LPS from *E. coli* and *S. minnesota*, we hypothesized that the LPS chemotypes differentially activate MyD88-dependent and -independent pathways. We used the MyD88 inhibitor ST2825, a synthetic analogue of MyD88 and a specific inhibitor of MyD88 dimerization, a crucial step for pathway activation^44^ (Fig. 6a). ST2825 completely abolished the signal induced by *E. coli* LPS in TLR4-HEK293 cells during the recording period, except for a negative small peak at the beginning (Fig. 6b). After stimulation with LPS from *S. minnesota*, ST2825 substantially reduced the signal until 2,400 s, however, the signal gradually increased and remained at around 60% compared to cells stimulated with LPS alone (Fig. 6b). These findings reveal that LPS *E. coli* and *S. minnesota* differentially activate MyD88-dependent pathways.

**Figure 6:**
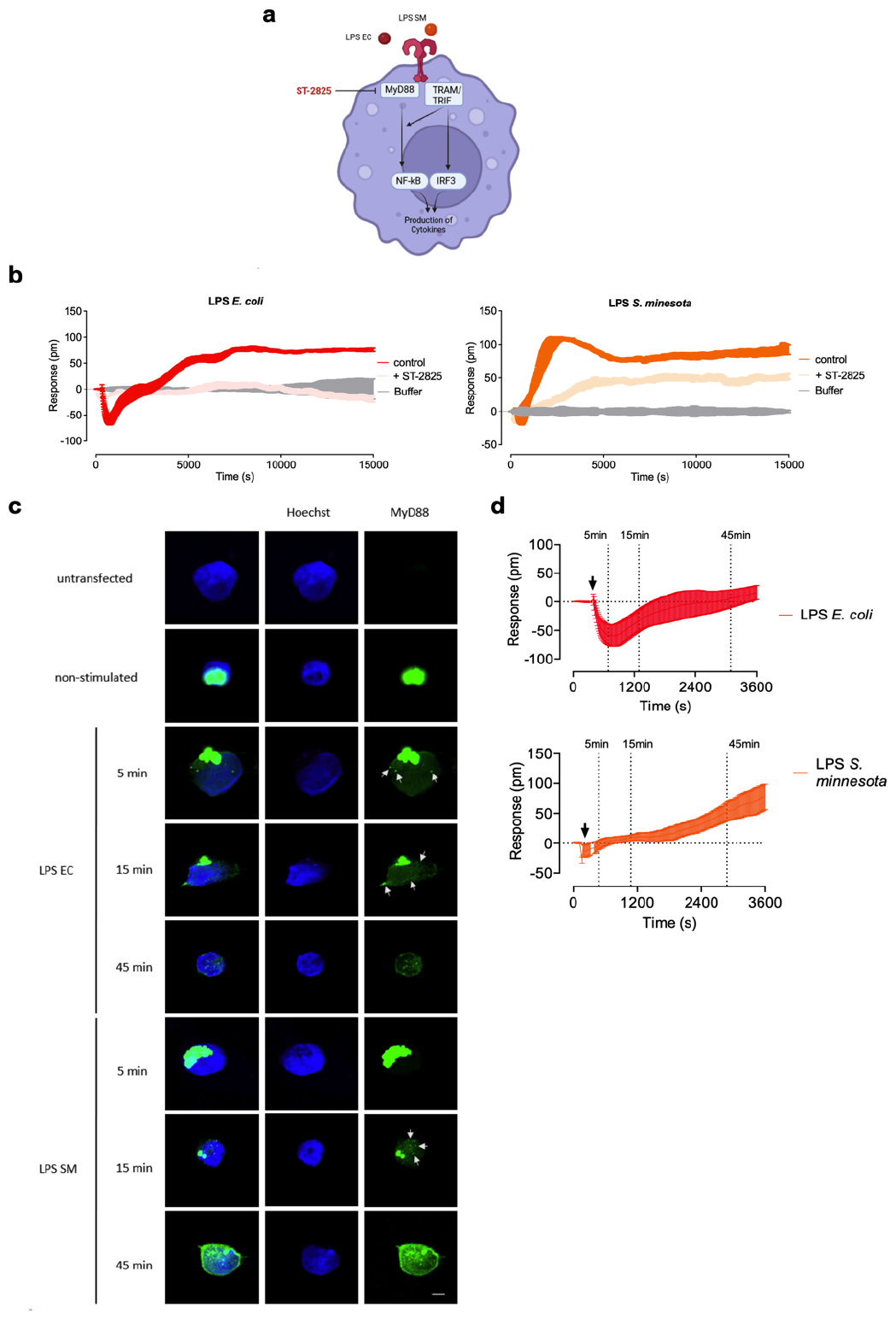
The use of the MyD88-inhibitor confirmed differences in signaling induced by different LPS sources. **(a)** Schematic representation of TLR4 signaling, ligands used and contact point of the MyD88 inhibitor ST2825. **(b)** Baseline corrected DMR representative recordings of HEK293 TLR4 reporter cells stimulated with LPS from *E. coli* (100 ng/ml) (b) or *S. minnesota* (100 ng/ml) or preincubated with 10 *μ*M of the MyD88 inhibitor ST2825. Shown are representative data of at least three independent experiments. **(c)** Immunofluorescence microscopy for localization experiments MyD88 (green) in transfected HEK Blue hTLR4 cells before and after stimulation with LPS *E. coli* and LPS *S. minnesota* (100 ng/ml) for 5 min, 15 min or 45 min. Cells are transfected with 200 ng MyD88-Venus construct and counterstained with the nuclear probe Hoechst (blue). Scale bars, 5 *μ*m. **(d)** Baseline corrected DMR representative recordings of HEK293 TLR4 reporter cells stimulated with LPS from *E. coli* (100 ng/ml) or *S. minnesota* (100 ng/ml). Shown are representative data of at least three independent experiments.

To uncover potential differences in the recruitment of the adapter protein MyD88 by LPS *E. coli* and *S. minnesota*, we transfected TLR4-HEK293 cells with Venus-tagged MyD88 and assessed cellular localization of MyD88 by immunofluorescence analysis. In the absence of LPS, MyD88 was located in condensed form in the cytoplasm as demonstrated previously^45^ (Fig. 6c). After stimulation with LPS *E. coli*, MyD88-Venus readily started to scatter from the compact structures and formed puncta already after 5 minutes, while in cells incubated with LPS S. minnesota, a delayed formation of MyD88-Venus puncta was observed (Fig. 6c, white arrows). After 45 minutes, the formation of puncta seemed complete for both LPS chemotypes. The kinetic differences in early MyD88 assembly by LPS *E. coli* and *S. minnesota* might be reflected in the signal traces observed by the optical biosensor (Fig. 6d).

### Optical biosensor assay decodes signaling of endosomal TLRs

In contrast to other TLRs that are typically located on the cell surface and identify bacterial danger signals, a subset of TLRs, including TLR3 and TLR8, are located within the cell endosomes (Fig. 7a). We investigated whether endosomal TLR signaling could also be detected using optical biosensor technology. HEK293 cells express endogenous TLR3^46^ and the selective ligand poly(A:U)^47^ induced a concentrationdependent negative response (Fig. 7b). High concentrations (100 and 250 *μ*g/ml) did not return to baseline during the 4 h recording period. To further characterize the concentrationdependent response of poly(A:U), concentration-effect curves calculated from the area under the curve were generated and an EC_50_ value of 45 *μ*g/ml was determined (Fig. 7c). In the absence of selective TLR3 inhibitors, we used the antimalarial drug and endosomal TLR inhibitor chloroquine^48^ to confirm the poly(A:U)-induced signal. Chloroquine alone showed no significant effect (Supplementary Fig. 6a) and clearly blunted the poly(A:U)induced signal (Fig. 7d).

**Figure 7:**
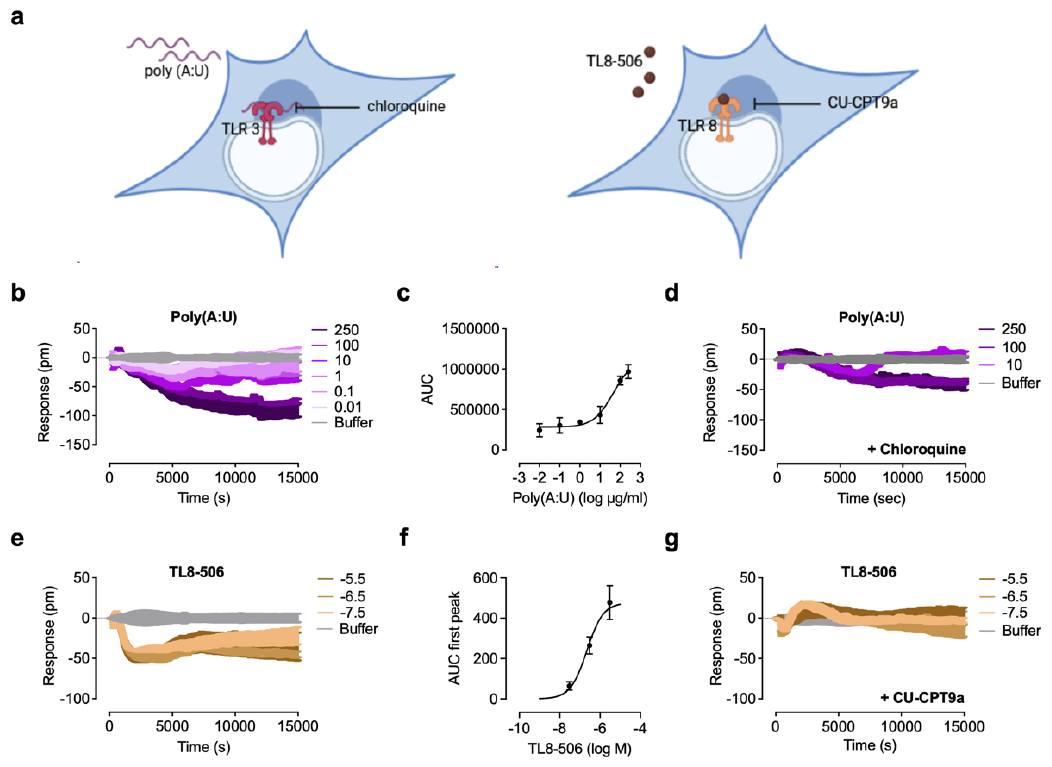
Optical biosensor decodes signaling of endosomal TLRs. **(a)** Schematic representation of TLR3 and TLR8 signaling, ligands used and contact point of the inhibitors chloroquine and CU-CPT9a. **(b)** Baseline corrected DMR representative recordings of HEK293 control reporter cells stimulated with the indicated concentrations (*μ*g/ml) of the TLR3 agonist poly(A:U). **(c)** Sigmoidal concentration effect curves resulting from DMR traces of at least three independent experiments (Mean ± S.E.M). Concentration-effect curves of DMR data were generated by the area under the curve (AUC). **(d)** Baseline corrected DMR representative recordings of HEK293 control reporter cells preincubated with 15 *μ*M chloroquine and stimulated with the indicated concentrations (*μ*g/ml) of poly(A:U). **(e)** Baseline corrected DMR representative recordings of HEK293 TLR8 reporter cells stimulated with the indicated concentrations of the TLR8 agonist TL8-506. **(f)** Sigmoidal concentration effect curves resulting from DMR traces of at least three independent experiments (Mean ± S.E.M). Concentration-effect curves of DMR data were generated by the area under the curve (AUC). **(g)** Baseline corrected DMR representative recordings of HEK293 TLR8 reporter cells preincubated with 1 *μ*M CU-CPT9a and stimulated with the indicated concentrations of TL8-506. Shown are representative data of at least three independent experiments.

TLR8 senses viral and bacterial uridin-rich singlestranded RNA within the endosomal compartment^49–52^ (Fig. 7a). The potent and selective TLR8 agonist TL8-506 triggered also a negative DMR signal in TLR8-transfected HEK293 cells with a plateau at 1,500 s (Fig. 7e). A concentration-effect curve calculated from the area under the curve of DMR traces showed an EC_50_ value of 0.2 *μ*M (Fig. 7f) which is in line with the EC_50_ of 0.6 *μ*M generated in a secreted embryonic alkaline phosphatase reporter assay^53^. CU-CPT9a, a highly potent and selective TLR8 inhibitor^54,55^ completely inhibited the TL8-506-induced signals (Fig. 7g) without producing a DMR shift (Supplementary Fig. 6b) indicating that the signal is TLR8 receptor-specific. No signals were detected in the presence of the TLR8 agonist in HaCaT or control HEK293 cells lacking TLR8 (Supplementary Figs. 6c and d). The observation that both TLR3 and TLR8 induce negative traces suggests that optical biosensors can distinguish between signal cascades induced by cell surface and endosomal TLRs.

## Discussion

Despite many efforts to decipher the precise signaling mechanisms of TLRs, large gaps remain in our understanding of TLR function and signaling^5^. The development of new TLR-based therapies has been partially hampered by incomplete knowledge of signaling complexities as exemplified by the failure of TLR4 antagonists in sepsis^56,57^. Receptor signaling, including TLRs, is often promiscuous and triggers several intracellular pathways after receptor activation by endogenous or exogenous ligands. Label-free DMR capturing morphological rearrangement enables real-time detection of integrated cellular responses in living cells and is a powerful tool for studying receptor activation and pathway signaling^6,33,36,58^. We are now able to visualize signals directly after TLR activation and uncover previously unrecognized mechanisms in TLR pharmacology. Optical biosensor technology identifies distinct spatiotemporal dynamics induced by ligands for cell surface and endosomal TLRs and cell type-dependent signaling even in primary cells. This technology also provides EC_50_ and E_max_ values, facilitating the determination of drug potency and efficacy.

Dimerization of TLRs triggers the assembly of supramolecular organizing centers in the cytosol, e.g. the myddosome and triffosome, leading to inflammatory and adaptive immune responses^5^. Although signaling cascades have been extensively studied, it is still unclear whether different TLR ligands targeting the same receptor show bias towards intracellular signaling pathways. Ligands can bias protein conformation through selective affinity, which differentially activate certain receptor signaling responses compared with others^13^. Ligand bias has been mainly studied for GPCRs^14,59–61^ and only very limited evidence is available for TLR4 and TLR9 ligands^62–64^. TLR4 activates MyD88-dependent and TRIF-dependent signaling pathways^41,42^. The ability to selectively target a single signaling pathway offers tools for fine-tuning of receptor signaling, and perhaps even receptor conformations. We show that different LPS chemotypes clearly diverge in DMR responses indicating biased signaling downstream of TLR4^65^. Our results are unequivocally in line with structure-activity relationship studies that show that structural differences in lipid A can result in varying degrees of inflammation. Lipid A from *E. coli* maximally stimulates the inflammatory response whereas lipid A from *S. minnesota* induces a lower inflammatory response through NF-κB and a greater stimulatory effect on IRF3 signaling^66–68^. These findings indicate that structural differences not only influence potencies but also result in biased agonism^4^. Unlike GPCR signaling, it might be challenging to precisely pinpoint a signal trace to a specific TLR downstream pathway, we were able to uncover kinetic differences of LPS chemotypes in MyD88 activation. The dynamics of cellular responses including activation of NF-kB, a major transcription factor involved in TLR signaling, is cell type dependent^69^. The different DMR signals obtained in epithelial, epidermal and immune cells induced by TLR4 and TLR2 ligands, respectively, underline the unique spatiotemporal dynamics. While it is widely accepted that TLR4 and TLR2 recognize different cell wall components during bacterial infection and share the same downstream NF-kB signal processing apparatus^70,71^, our findings reveal that inducing the same signal partner does not necessarily result in the same signaling pathway dynamics or kinetics. When stimulated independently by specific TLR4 and TLR2 agonists, distinct dynamic NF-κB profiles appeared in single cells^72^. These findings are consistent with signal traces detected with optical biosensor assays, which also allow for the characterization of NF-κB dynamics induced by TLR ligands.

Furthermore, optical biosensor technology uncovers differential TLR2 signaling between monocytes and macrophages^73^ during a single experiment. These results suggest that activation of TLR heterodimers by the different agonists induce different signaling cascades/kinetics in monocytes and macrophages and provide evidence for ligand- and system-specific bias which has been reported for GPCRs^74^. Real-time analysis also provides new opportunities to visualize specific signaling patterns in cells that are equipped with different signaling molecules.

The optical biosensor assay is also sensitive enough to detect ligand-specific optical traces, as demonstrated for synthetic lipopeptides with Pam_3_CSK_4_ displaying delayed activation kinetics at TLR2/1 compared to Pam_2_CSK_4_ when activating TLR2/6^75^. Notably, Pam_3_CSK_4_ and Pam_2_CSK_4_ lead to distinct signatures after TLR2/1 activation suggesting that TLR2 heterodimers induce ligand-dependent biased signaling. This finding has remained elusive using conventional end-point assays but uncovered by optical biosensor technology.

TLR3, TLR7, TLR8, TLR9 are expressed intracellularly within the endoplasmic reticulum, endosomes and lysosomes^76^. A unique feature of TLR3 is the ability to signal independently of the adaptor protein MyD88^77^. Activation of the receptor leads to the recruitment of the adaptor TRIF which associates subsequently with TRAF3 and TRAF6. Further steps activate both NF-κB, MAPKs^78,79^ and IRF3^80^. Given the evidence that chloroquine reduces TLR3/IRF3/IFN-β signaling^81^ that the residual signal after incubation with chloroquine reflects a pathway that is still active. Different signal traces following TLR3 and TLR8 activation may also provide evidence for the involvement of different adaptor proteins, e.g., MyD88. It is likely that biased signaling is initiated at endosomal TLRs by nucleic acids of varying origins to fine-tune innate immune responses.

Biosensor technology can reveal the complexity of signaling pathways, where numerous mediators contribute to optical signatures. Yet, it is important to note that labelfree responses are closely linked to a ‘black box’ technology. When combined with pharmacological pathway inhibitors or powerful genome editing tools such as CRISPR/Cas9 technology, specific proteins involved in TLR signaling as reflected by DMR signals can be identified thus complementing traditional experimental assays. Our approach is also an option to investigate other pattern recognition receptors such as NOD-like receptors and RIG-I-like receptors and TLR agonists or antagonists on patient tissues, reflecting disease relevant conditions^82,83^. By using optical biosensor technology differences in signaling pathway activation in the tissue removed from the patient could be immediately recognized. Together with the possibility to monitor the cellular responses in real-time this method could give indications of illnesses or changes and thus further deepens the understanding of dysregulated TLR signaling.

Real-time analysis of receptor dynamics provides a comprehensive understanding of events that occur earlier than what is typically observed with assays that report an effect at a specific endpoint or signaling step at several fixed time points. Assays based on optical biosensor technology allow for the detection of signaling events that have been previously overlooked and readily identify and characterize ligands that act on the same receptor but differentially activate downstream pathways. Our findings offer an unprecedented approach and warrant further investigation of the dynamics of Toll-like receptor signaling to foster the development and characterization of new Toll-like receptor modulators.

## Acknowledgements

This work was funded partially by the German Research Foundation to G.W. (grant RA 895/16-1). We thank Dr. Nicole Merten for technical support. We would like to thank the Next Generation Sequencing Core Facility (https://btc.uni-bonn.de/dfg/) of the Medical Faculty at the University of Bonn for providing support and instrumentation funded by the Deutsche Forschungsgemeinschaft (DFG, German Research Foundation). Figs. 1a, 1b, 2a, 6a, and 7a were created with Biorender.com. This paper was typeset with the bioRxiv word template by @Chrelli: www.github.com/chrelli/bio-Rxiv-word-template

## Author contributions

J.H. and F.L. performed experiments; J.H., F.L., and G.W. analyzed the data; E.K. provided reagents; G.W. directed the study; J.H., F.L., E.K., and G.W. wrote the manuscript.

## Materials and Methods

### Ethics statement

The studies with human blood were approved by the ethics committee of the University Clinic Bonn (315/22) and written informed consent was obtained from all healthy donors.

### Chemical compounds

The TLR ligands Pam_3_CSK_4_ and Pam_2_CSK_4_ (synthetic triacylated and diacylated lipoproteins), lipopolysaccharide from *Escherichia coli* O111:B4 (LPS *E. coli*), *Salmonella minnesota* R565 (LPS *S. minnesota*) and *Rhodobacter sphaeroides* ultrapure, TL8-506, CU-CPT9a as well as poly(A:U) were purchased from InvivoGen (Toulouse, France). TAK-242 was from BioTechne GmbH (Wiesbaden, Germany). Acetylcholine iodide and atropine sulfate, epinephrine, forskolin, chloroquine-diphosphate as well as the TLR2 antagonist CU-CPT22 were purchased from Sigma Aldrich Chemie (Steinheim, Germany). Iperoxo was synthesized as described previously^84,85^. The TLR2 antagonist C29 was obtained from ChemDiv (San Diego, USA). The inhibitor ST2825 (MyD88) was obtained from MedChemExpress (New Jersey, USA).

### Cloning of MyD88-Venus construct

MyD88 was synthetized and inserted into the HindIII and XhoL restriction sites of vector plasmid pcDNA-3.1 by Genecust (Boynes, France). Venus-Ggamma-pcDNA3.1 was a gift from Kirill Martemyanov. cDNA encoding human MyD88, including a N-terminally HindIII restriction site and a C-terminally GGATCC linker, was amplified by PCR (primers: GTCAGCAAGCTTATGGCCGCTGGCGGACCTGG, GCTCCTCGCCCTT GCTCACCATGGATCCACCTCCGGGCAGGCTCAGGGCTTTGGC). XhoI was inserted C-terminally into the cDNA of Venus using amplification by PCR (primers: ATGGTGAGCAAGGGCGAGGAGC, GCTGACCTCGAG TTACTTGTACAGCTCGTCCATGCCGAG). The plasmid expressing C-terminally Venus-tagged MyD88 (MyD88-Venus) was generated by overlap-extension PCR of the amplified MyD88 and Venus coding sequences (primers: GTCAGCAAGCTTATGGCCGCTGGCGGACCTGG, GCTGACCTCGAGTTACTTGTACAGCTCGTCCATGCCGAG) and subcloning into the HindII and XhoI sites of pcDNA3.1.

### Cell culture and primary cells

HEK Blue hTLR2-TLR1, hTLR2-TLR6, hTLR2KO-TLR1/6, hTLR4, hTLR8 and Null2 cells (InvivoGen, Touluse, France) were cultured in DMEM medium supplemented with 10% fetal bovine serum (FBS), penicillin (100 U/ml), streptomycin (100 *μ*g/ml), L-glutamine (2 mM), normocin (100 *μ*g/ml) and selective antibiotics (selection: hTLR2/1, hTLR2/6, hTLR4; zeocin (100 *μ*g/ml): hTLR8, Null2; blasticidin (30 *μ*g/ml): hTLR8). HEK-293 cells were grown in DMEM high glucose supplemented with 10% FCS, l-glutamine (2 mM), 100 U/ml penicillin, and 100 *μ*g/mL streptomycin and HaCaT cells were cultured in RPMI 1640 supplemented with 10% FCS, l-glutamine (2 mM), 100 U/ml penicillin, and 100 *μ*g/mL streptomycin. THP-1 cells (passage 4-25) (DSMZ, Braunschweig, Germany) were cultured as previously shown^86,87^. Cells were kept at 37 °C in a humidified atmosphere with 5% CO_2_. HEK-Blue Null2 and hTLR2KO-TLR1/6 cells were used as control as previously described^30,88,89^.

PBMCs (peripheral blood mononuclear cells) were isolated from buffy-coat donations (Institute of Experimental Haematology and Transfusion Medicine, University Clinic Bonn) by density gradient centrifugation as described previously^90^ using Biocoll separation media (Bio&Sell, Nuremberg, Germany). PBMCs were washed three times with PBS containing EDTA. After the third washing step, 5x10^6^ (24 well) or 70.000 (384 well) PBMCs were seeded per well to isolate the contained primary monocytes due to plastic adhesion. After 1 h incubation at 37 °C in a 5% CO_2_ incubator the remaining suspension cells were removed from the adherent primary monocytes by washing the cells three times with PBS. Before stimulation, the cells were incubated in media with 10% FCS and antibiotics (penicillin/streptomycin) for 24 h at 37 °C in a 5% CO_2_ incubator.

### Dynamic mass redistribution label-free assay

DMR assays were performed using the EPIC system (Corning) according to established protocols^33,36,91^. Briefly, the day before the assay, cells were seeded at 20,000 cells/well (HEK293 cells), 10,000 cells/well (HaCaT cells) into either an Epic 384-well uncoated or fibronectin-coated glass microplate (Corning, New York, NY, USA) and cultured for 24 h to obtain confluent monolayers. For generation of macrophages, THP-1 monocytes were seeded (20,000 cells/well) in medium containing 25 ng/ml phorbol 12-myristate 13-acetate (PMA, Sigma-Aldrich) for 48 h and afterwards rested for 24 h. In the case of THP-1 monocytes, cells were seeded as suspension cells in assay buffer on the day the experiment was performed. At the day of the assay, cells were washed twice with 50 *μ*l assay buffer (Hank’s Balanced Salt Solution (HBSS) containing 20 mM HEPES (pH 7.0) and centrifuged for 10 seconds to ensure that no drops adhered to the sides of the well, and that cells were in contact with the bottom (biosensor). The final volume in each well was 30 *μ*l. Epic microplates were incubated post cell-seeding for 1.5 hours in the EPIC instrument at 37 °C. Serial compound dilutions were made in the same assay buffer. For DMR measurements the Epic biosensor (Corning) was used. After reading baseline, compounds were added using a semiautomatic liquid handler Selma (Analytik Jena AG, Jena, DE). The addition of 10 *μ*l compounds were carried out in a volume of 30 *μ*l/well. DMR signals were recorded for 15,000 seconds, and data analyzed and exported with the Epic Analyzer Software (Corning). All DMR signals were baseline corrected. Antagonists or inhibitors were incubated 2h before agonist injection. Compound responses represented in traces were described as picometer (pm) shifts over time (s) following baseline normalization. All the experiments were carried out at 37°C. For a detailed description of the methods see^36^. DMR experiments were carried out in triplicates or quadruplicates.

### ELISA

Cell culture supernatants were collected and analyzed for IL-8 release using a commercially available ELISA kit (Thermo Fisher Scientific).

### LDA assay

LDH assay was performed according to the manufacturer’s instructions (CyQUANT LDH Cytotoxicity Assay, Thermo Fisher Scientific). The percentage of LDH release was calculated compared to 100% cell lysis control.

### Immunofluorescence

HEK Blue hTLR4 cells (Invivogen, Touluse, France) were seeded in a density of 0.85x10^6^ cells per well in 6-well plates in media containing 10% FCS and antibiotics (penicillin/streptomycin/normocin/selection). After 24 h incubation at 37 °C in a 5% CO_2_ incubator, the media was changed to media with 5% FCS with antibiotics. Subsequently, the cells were transiently transfected with 200 ng MyD88-Venus construct using PEI Max (Polysciences) according to manufacturer’s protocol and were incubated for further 24 h at 37 °C in a 5% CO_2_ incubator. The transfected cells were seeded in L-polylysin (Poly-L-lysin hydrobromid, P6282, Sigma-Aldrich) coated 8 well chamber slides in a total of 7x10^4^ cells per well and were incubated for 48 h at 37 °C in a 5% CO_2_ incubator. For stimulation the cells were incubated with LPS *E. coli* (100 ng/ml) or LPS *S. minnesota* (100 ng/ml) for 5 min, 15 min or 45 min. After stimulation the cells were fixed using 4% paraformaldehyde (Carl Roth) for 10 min. Between each staining step the cells were washed extensively using PBS. The cells were permeabilized with 0.2% Triton X-100 (Sigma) for 10 min and blocked for 30 min using 10% normal goat serum (Cell Signaling Technology, 5425). The nucleus was stained using Hoechst 33342 (1 *μ*g/ml, Thermo Fisher Scientific, 62249) and the cells were mounted in ProLong Glass Antifade Mountant (Invitrogen, P36982). Images were taken with the confocal laser microscope Nikon Ti-E with A1 confocal scanner and were processed using NIS-Elements Viewer 5.21.

### RNA-seq

Total RNA was harvested according to the manufacturer’s instructions (innuPREP RNA Mini Kit 2.0, Analytik Jena). RNA libraries were prepared using QuantSeq 3’-mRNA Library Prep Kit (Lexogen) and RNA-seq was performed with the NovaSeq 6000 (Illumina).

### RNA-seq data analysis

The nf-core rnaseq pipeline^92^ (version 3.10.1) was applied for the preprocessing and the quantification of the reads using default parameters. Here the first step is quality and adapter trimming with TrimGalore (version 0.6.7). This will be followed by aligning the trimmed reads against human genome (GRCh38) with STAR^93^ (version 2.7.9a). The aligned data are then used as input for Salmon^94^ (version 1.9.0), which employs pseudoalignment to estimate transcript abundances. The transcript-level quantifications will then be aggregated to obtain gene-level expression estimates.

The statistical analysis was executed in the R environment (version 4.2.0, cite R). Given the notable biological variability observed between the two cell lines, HEK and THP, distinct analyses were undertaken for each cell line. To ensure the robustness of the results, only genes with a minimum count of 10 in at least two samples were utilized for the inference analysis. The Bioconductor package DESeq2^95,96^ was employed for the differential gene expression analysis. Subsequently, the Benjamini-Hochberg method was applied to calculate multiple testing-adjusted p-values (false discovery rate, FDR) for each contrast. For visualization, the packages ggplot2^97^ was used to generate volcano plots.

### Data and statistical analysis

Data are shown as means ± S.E.M. from at least three independent experiments and data points from individual experiments. DMR experiments were depicted with one representative trace. To compare two means, statistical significance was based on a Student’s t-test with P < 0.05. Comparisons of groups were performed using one-way-ANOVA analysis with a Tukey-Kramer Post-Test. In all cases, P < 0.05 (*), was considered as the level of statistical significance, P < 0.01 (**) and P < 0.001 (***) was indicated for selected comparisons.

All experimental data were analyzed by using Graph Pad Prism 8.0 (GraphPad Software Inc., San Diego, CA, USA). Data obtained from DMR experiments were analyzed by a four-parameter-logistic function yielding parameter values for a ligand’s potency (logEC_50_) and maximum effect (E_max_). For data normalization, indicated as relative response (%), top levels of concentration effect curves at 15,000 s of the data set were set 100% and bottom levels 0%.

**Supplementary Figure 1:**
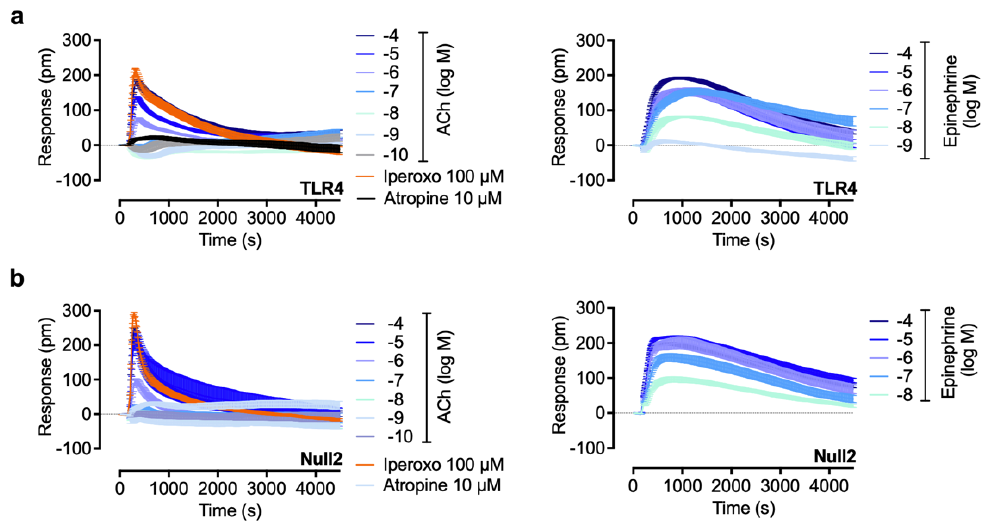
**(a,b)** HEK293 TLR4 reporter cells **(a)** or control HEK293 cells **(b)** stimulated with the indicated concentrations of the acetylcholine (ACh) and epinephrine. ACh visualizes signaling along the Gq_α/11_ pathway whereas epinephrine induces presumably the G_s_ pathway. This illustrates that cells overexpressing TLR4 show consistent GPCR signaling patterns when compared to control cells lacking TLR4. DMR recordings are representative (mean ± S.E.M.) of at least three independent biological replicates.

**Supplementary Figure 2:**
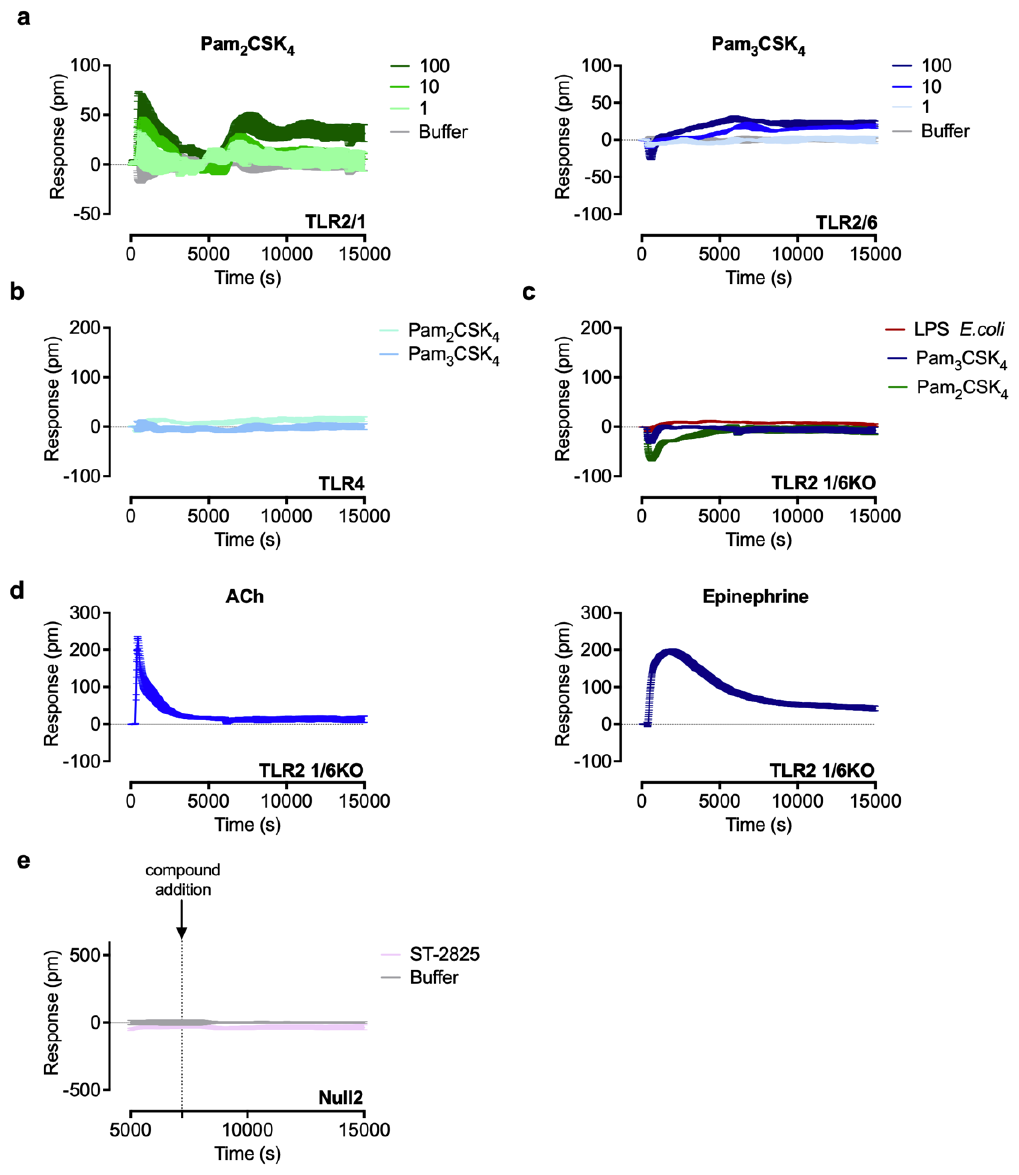
**(a)** HEK293 TLR2/1 or 2/6 reporter cells stimulated with the indicated concentrations (ng/ml) of the TLR2 agonists Pam_2_CSK_4_ (TLR2/6) and Pam_3_CSK_4_ (TLR2/1). **(b)** HEK293 TLR4 reporter cells stimulated with the TLR2 agonists Pam_2_CSK_4_ (100 ng/ml) and Pam_3_CSK_4_ (100 ng/ml). **(c,d)** HEK293 TLR21/6 knock-out cells stimulated with **(c)** the TLR4 agonist LPS *E. coli* (10 ng/ml), TLR2/1 agonist Pam_3_CSK_4_ (100 ng/ml), TLR2/6 agonist Pam_2_CSK_4_ (100 ng/ml), and **(d)** acetylcholine (100 *μ*M) or epinephrine (100 *μ*M). **(e)** HEK293 control cells lacking TLR4 (Null2) incubated with the MyD88 inhibitor ST2825 (10 *μ*M). DMR recordings are representative (mean ± S.E.M.) of at least three independent biological replicates.

**Supplementary Figure 3:**
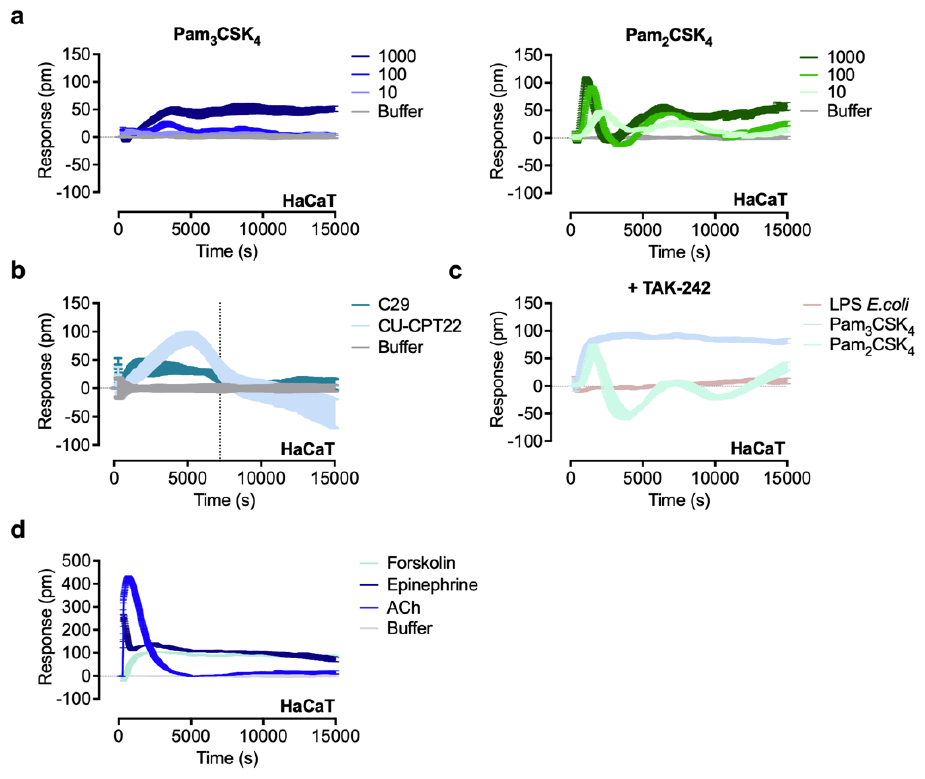
**(a)** HaCaT cells stimulated with the indicated concentrations (ng/ml) of TLR2/1 agonist Pam_3_CSK_4_ or TLR2/6 agonist Pam_2_CSK_4_. **(b)** HaCaT cells incubated with the TLR2 antagonists C29 (50 *μ*M) and CU-CPT22 (50 *μ*M). The depicted line visualizes the timepoint 2 h. This time corresponds to the preincubation time of the inhibitors. Since the signal is back at the baseline additive effects are not suspected. **(c)** HaCaT cells stimulated with the TLR4 agonist LPS *E. coli* (100 ng/ml), TLR2/1 agonist Pam_3_CSK_4_ (100 ng/ml) or TLR2/6 agonist Pam_2_CSK_4_ (100 ng/ml) preincubated with the TLR4 inhibitor TAK-242 (50 *μ*M). **(d)** HaCaT cells stimulated with ACh (100 *μ*M), epinephrine (100 *μ*M) and forskolin (10 *μ*M). DMR recordings are representative (mean ± S.E.M.) of at least three independent biological replicates.

**Supplementary Figure 4:**
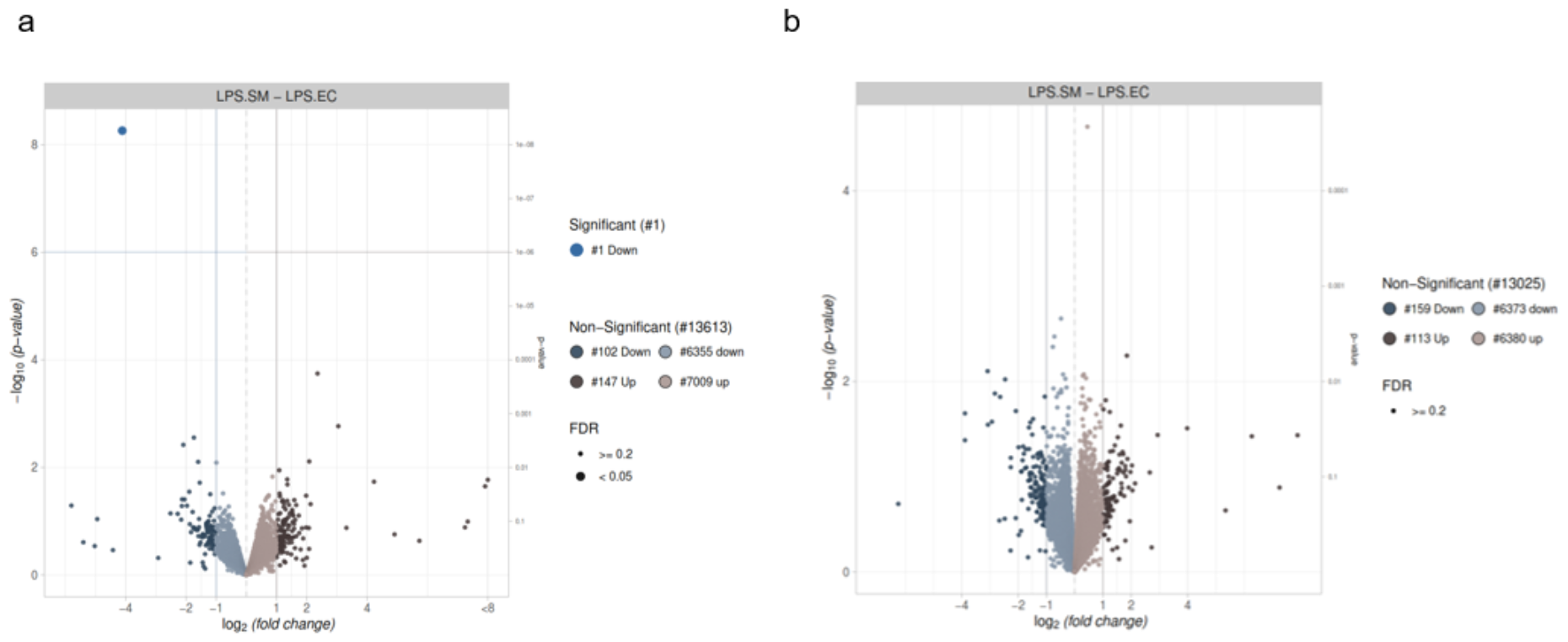
**(a, b)** The volcano plots of LPS *S. minnesota* vs. LPS *E. coli* in **(a)** HEK293-TLR4 cells or **(b)** THP-1 macrophages. For visualization, the package ggplot2 was used.

**Supplementary Figure 5:**
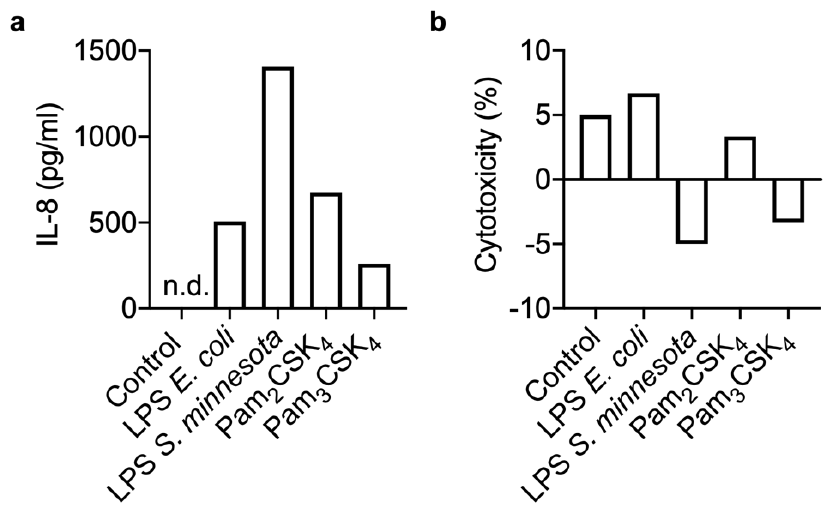
**(a)** IL-8 secretion from primary monocytes or **(b)** cytotoxicity effect following treatment with LPS *E. coli* (10 ng/ml), LPS *S. minnesota* (10 ng/ml), Pam_2_CSK_4_ (1 *μ*g/ml) or Pam_3_CSK_4_ (1 *μ*g/ml) for 4 h. IL-8 concentration in the cell-free supernatant was determined by ELISA. Data show results of one blood donor.

**Supplementary Figure 6:**
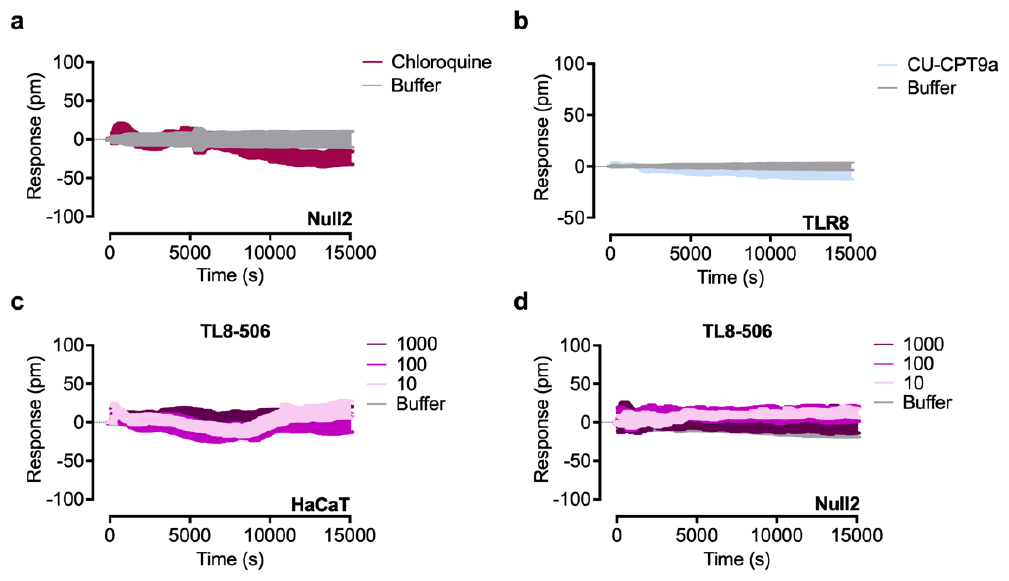
**(a)** Control HEK293 cells stimulated with chloroquine (15 *μ*M). **(b)** HEK293 TLR8 reporter cells stimulated with the TLR8 inhibitor CU-CPT9a (1 *μ*M). **(c)** HaCaT and **(d)** control HEK293 cells stimulated with the indicated concentrations (ng/ml) of the TLR8 agonist TL8-506. DMR recordings are representative (Mean ± S.E.M.) of at least three independent biological replicates.

